# A coarse-grained methodology identifies intrinsic mechanisms that dissociate interacting protein pairs

**DOI:** 10.1101/2020.07.10.196717

**Authors:** Haleh Abdizadeh, Farzaneh Jalalypour, Ali Rana Atilgan, Canan Atilgan

## Abstract

We address the problem of triggering dissociation events between proteins that have formed a complex. We have collected a set of 25 non-redundant, functionally diverse protein complexes having high-resolution three-dimensional structures in both the unbound and bound forms. We unify elastic network models with perturbation response scanning (PRS) methodology as an efficient approach for predicting residues that have the propensity to trigger dissociation of an interacting protein pair, using the three-dimensional structures of the bound and unbound proteins as input. PRS reveals that while for a group of protein pairs, residues involved in the conformational shifts are confined to regions with large motions, there are others where they originate from parts of the protein unaffected structurally by binding. Strikingly, only a few of the complexes have interface residues responsible for dissociation. We find two main modes of response: In one mode, remote control of disassociation in which disruption of the electrostatic potential distribution along protein surfaces play the major role; in the alternative mode, mechanical control of dissociation by remote residues prevail. In the former, dissociation is triggered by changes in the local environment of the protein e.g. pH or ionic strength, while in the latter, specific perturbations arriving at the controlling residues, e.g. via binding to a third interacting partner is required for decomplexation. We resolve the observations by relying on an electromechanical coupling model which reduces to the usual elastic network result in the limit of the lack of coupling. We validate the approach by illustrating the biological significance of top residues selected by PRS on select cases where we show that the residues whose perturbation leads to the observed conformational changes correspond to either functionally important or highly conserved residues in the complex.

## 1. Introduction

Chemical and physical processes within assemblies of proteins in the cellular environment are events often encompassing multiple time and length scales. Therefore, different modeling tools are commonly used to describe the network of interactions featuring the protein dynamics (Adcock and McCammon, 2006; Tozzini, 2010). In this vein, coarse-grained models (CG), with several atoms of the protein grouped into one bead and in the absence of atomic details of the solvent molecules have been developed to supplement the extremely expensive atomistic modeling of large scale motions of biomolecular aggregates (Orozco, 2014; Atilgan, 2018). CG models have proved to aid sampling efficiency, predict allosteric regulations (Ming and Wall, 2005) and describe conformational transition pathways (Kim et al., 2002; Miyashita et al., 2003). One useful measure of large-scale protein mechanics in the context of CG models is the elastic network model (ENM) (Tirion, 1996; Bahar et al., 1997; Hinsen, 1998). ENMs are based on the assumption that the potential energy of the system is approximated harmonically about a single minimum energy conformation. Methodologically, in ENM, the atomic details of the biomolecule structure are reduced to a network of nodes (typically one site per residue) connected by harmonic springs. Since all springs are in a relaxed state in the network, no energy minimization is required, in comparison to normal mode analysis in which an expensive initial energy minimization is required prior to calculating the Hessian matrix. For large biomolecules and multi-protein complexes, ENM models with a resolution lower than standard have been used (Durand et al., 1994; Doruker et al., 2002; Chennubhotla et al., 2005; Ahmed and Gohlke, 2006; Kurkcuoglu et al., 2009; Ross et al., 2018a). The gross representation of large assemblies has proven to predict dynamics of the rigid and flexible parts of the proteins (Ross et al., 2018b). Anisotropic network model (ANM) and Gaussian network model (GNM) are amongst the most widely used ENM-based methods (Bahar et al., 1997; Atilgan et al., 2001). While GNM is applied to produce the amplitudes of isotropic thermal motions, ANM describes both amplitudes and directionality of anisotropic motions. Increased amount of data for proteins of different forms (free, liganded, or complexed), elucidates the correlation between protein function observed in experiments and the global motions predicted by ANM/GNM analyses. Numerous studies have employed ENM-based models to explore various aspects of protein structural dynamics. These include identifying and visualizing collective motions (Kong et al., 2003), predicting modes of motion underlying function (Baysal and Atilgan, 2001b; Keskin et al., 2002; Zheng and Doniach, 2003; Whitford et al., 2007), and explaining details of conformational changes of various types and amplitudes (Tama and Sanejouand, 2001; Krebs et al., 2002; Zheng et al., 2007). ENMs may be applied to the refinement of medium to low-resolution structures of electron microscopy density maps of large macromolecular complexes or molecular envelopes derived from small-angle x-ray scattering (SAXS) data (Delarue and Dumas, 2004; Hinsen et al., 2005). Within the concept of ENM, researchers have developed approaches to generate feasible pathways for conformational transitions between two end conformers (Kim et al., 2002; Orellana et al., 2019), removing the need for expensive molecular dynamics (MD) simulations and all-atom empirical force fields to set up intermediate conformations. ENMs are also applied to determine the main evolutionary transformations of structural changes among homologous proteins (Leo-Macias et al., 2005; Han et al., 2008). In such an approach, for a given set of proteins, evolutionary direction is argued to be a combination of a small subspace projected by a few low frequency modes imposed by inter-residue contact topology.

We have extended ENM to analyze allosterically significant residues and function-related motions of proteins via a technique named perturbation-response scanning (PRS) (Atilgan and Atilgan, 2009; Atilgan et al., 2010a). The methodology inserts fictitious forces on selected atoms and predicts the response within the realm of linear response theory (LRT). In vivo, the perturbation may arrive in the form of changing environmental conditions such as pH or ionic strength (Abdizadeh et al., 2015a; Sensoy et al., 2017), or it may act directly on the chain as in pulling (Carrion-Vazquez et al., 2003; Dietz et al., 2006) or other single-molecule experiments, as well as through mutations or ligand binding. PRS serves as a tool to gain insight into molecular origins of mechanical feedbacks of bimolecular structures through recording response to each inserted force on each residue of a protein (Atilgan and Atilgan, 2009; Atilgan et al., 2010a; Atilgan et al., 2011; Abdizadeh et al., 2015b; Abdizadeh and Atilgan, 2016). It is further capable of recognizing how directionality of the inserted force may coordinate the response of the protein in a functional motion (Jalalypour et al., 2020). PRS requires two distinct conformations of a protein, determined, e.g. by x-ray crystallography, as input; and relies on LRT to relate virtual external forces acting on a protein to the perturbed positions of the residues (Ikeguchi et al., 2005). In PRS, one scans a protein structure residue by residue through applying forces in many directions and records the subset of responses that encourage conversion to another known conformation of the protein. Thus, one can map the regions on the protein surface whose perturbation might lead to the expected conformational change. Besides mapping active site residues that are prime regions for invoking conformational transitions, this approach also has the potential of pointing out allosteric locations and drug target regions. For example, previously, we have studied the proteins calmodulin (Atilgan et al., 2011) and ferric binding protein (Atilgan and Atilgan, 2009) via PRS. By mutating those residues that were implicated in allosteric communication, we later verified through classical MD simulations that they affect the conformation distributions (Aykut et al., 2013; Guven et al., 2014). In a later study, we have performed PRS on subtilisin in complex with its inhibitor to pinpoint hot residues involved in catalytic mechanism and stability of the enzyme (Abdizadeh et al., 2015b). PRS has also been used in the conformation generation step of a flexible docking scheme for exploring protein-ligand interactions (Bolia et al., 2014). In a similar methodology, Mottonen et al. have used a method based on distance constraint model to impose constraints on the torsional degrees of freedom of the protein to mimic a hypothetical ligand-binding situation (Mottonen et al., 2010; Jacobs et al., 2012).

In this manuscript, we utilize these CG approaches to address a major challenge for structural biology in providing a mechanistic view of the behavior of molecular complexes and their conformational changes. Protein-ligand and protein-protein interactions (PPI) govern most of the cellular processes (De Las Rivas and Fontanillo, 2010). Many studies investigate the protein-ligand complexes and look for functional regions, binding sites or druggable cavities (Lichtarge and Sowa, 2002; Xie et al., 2009; Siragusa et al., 2015). On the other hand, PPI allow a protein to perform its biological function by interacting with another partner protein. (Ozdemir et al., 2019) Therefore, the interface is usually considered as a candidate to be targeted by a potential drug such as orthosteric or allosteric PPI modulators (Xie et al., 2009; Ni et al., 2019). Studying PPIs and protein-interaction networks may provide insights into new opportunities in the medical, biotechnological, and pharmaceutical fields. Hence, several approaches have been proposed to study PPIs (Ozdemir et al., 2019). A number of bioinformatics techniques have been developed to predict PPI networks based on genomic-context, sequence homology and structural similarity (Shi et al., 2005). Most systematic studies involving protein-protein complexes focuses on the interaction interface to determine compatibility of the structures or attempts to study individual PPI and predicts residues, called hotspots, effective in recognition of partner proteins (Liu et al., 2018a; Qiao et al., 2018). Alanine scanning mutagenesis is the major experimental method to identify hotspots (Kenneth Morrow and Zhang, 2012). In one study, non-covalent interactions (hydrophobic, van der Waals, and hydrogen bonding) are found to account as the major forces operating at the PPI interfaces (Gao et al., 2004). Available computational techniques for PPI hotspot prediction are roughly divided to two groups whereby most use the complex structure and a few utilize unbound structures (Ozdemir et al., 2019). Generally, hotspots resulting from the computations are compared to those from alanine scanning mutagenesis experiments (Bradshaw et al., 2011; Ibarra et al., 2019). In addition, machine learning-based methods have been developed to predict hotspots, considering the amino-acid features and conservation information (Liu et al., 2018b). Most recently, by ignoring the internal structures of the molecules and scanning the protein surface for the so-called “interaction fingerprints,” geometric deep learning algorithms have been developed for predicting protein-protein complexes (Gainza et al., 2020). Although these approaches attempt to define a general interaction pattern based on parameters such as structure, hydrophobicity or polarity, there is no general rule to be used in PPI prediction due to their diversity (Ni et al., 2019).

In this work we address a reverse problem: How is it possible to trigger dissociation events between proteins which have already formed a complex? We study 25 sets of protein complexes utilizing PRS with the ENM harmonic potential to determine regions responsible for rendering known complexes incompatible. Elastic network construction helps one to probe conformational changes due to altered physical and chemical environment (Atilgan et al., 2012). Accordingly, the information needed for assessing protein-protein interactions can be derived from knowledge of inter-residue contact topology, buried in the Hessian matrix deduced from ENM (Bahar et al., 2010). Rather than focusing on the interface of the interacting subunits, we relate the physical effects of the internal dynamics of protein complexes to the motions involved in their disassociation. We show that PRS maps residues that may alone initiate the structural change between the bound and unbound forms during disassociation processes of the protein complexes.

## 2. Models and Methods

The conformational change was analyzed via PRS between two different conformers of a protein, one in its complexed form with another protein, and the other in its unbound form. The propensity to convert between conformations was examined for these two states of the protein by employing fictitious forces. The detailed theory of PRS has been laid-out in previous studies (Atilgan and Atilgan, 2009; Atilgan et al., 2010a). In brief, the unbound state of a protein may be described by a perturbation of the Hamiltonian of the bound state, or vice versa. Under LRT, the shift in the coordinates due to unbinding is approximated by (Yilmaz and Atilgan, 2000; Atilgan and Atilgan, 2009):

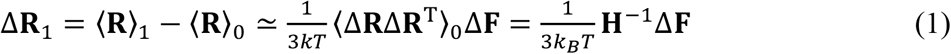

where the subscripts 1 and 0 denote perturbed and unperturbed configurations of the protein, *k_B_* is the Boltzmann constant and *T* is temperature. **ΔF** vector contains the components of the externally inserted force vectors on the selected residues; e.g. for the perturbation of a single residue *i*, 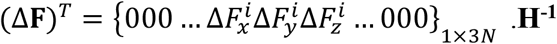 is the variance-covariance matrix which may be obtained from either the atomic coordinate trajectories of MD simulations of suitable length (Atilgan et al., 2012), or by imposing the approximation of harmonic springs between pairs of interacting atoms. In this work, we generate the **H^-1^** matrix from a coarse-grained approach, constructing a network of *N* nodes on the C_α_ atoms of the protein complexes whose coordinates are directly used from their protein data bank (PDB) entries. Any given pair of nodes interacts in accord with a conventional harmonic potential, if the nodes are within a cut-off distance, *r_c_*, of each other. This leads to a total of *M* interactions. Within the scope of an elastic network of residues that are connected to their neighbors by linear-elastic springs, one gets the 3*N×M* direction cosine matrix **B** (Yilmaz and Atilgan, 2000). **BB**^T^ is exactly the Hessian if harmonic interactions with uniform force constants for all *M* bonds in the network are assumed. (**BB**^T^)^-1^ is the covariance matrix **H^-1^** for a given configuration, which is also an *N×N* supermatrix whose *ij^th^* element is the 3×3 matrix of correlations between the x-, y-, and z-components of the fluctuations **ΔR**_i_ and **ΔR**_j_ of residues *i* and *j*.

**H** of the system has at least six zero eigen-values corresponding to the purely translational and rotational motions. The eigen-value distribution of the Hessian of proteins is such that the low-frequency region is more crowded than expected of polymers or other condensed matter (Ben-Avraham, 1993). Thus, the choice of the cutoff distance, *r_c_*, for the construction of the Hessian is critical for extracting protein-like properties from the systems studied (Atilgan et al., 2010b). For all the proteins studied in this work, we coarse-grain the crystal structure so that each residue is represented by the coordinates of its *C_α_* atom. To account for the flexibility of proteins, we repeat the PRS analysis for a variety of cut-off distances in the range of 10.0–14.0 Å in increments of 1 Å; the lower limit of 10 Å agrees with the definition of first coordination shell of residues in proteins (~ 7 Å). For each network structure, we ensure that the system has six zero eigen-values corresponding to the translational and rotational degrees of freedom of the protein. The smallest common *rc* at which we obtain six zero eigenvalues for all the proteins tested is 9 Å.

PRS technique relies on repeating the above LRT calculation (eq. 1) by scanning the residues of the protein one-by-one and focusing further on those perturbations that overlap with the conformational change, Δ**R**_1_ = <**R**>_1_ - <**R**>_0_. There is no a priori assumption on how a force might be generated at a particular point. Conversely, after finding the force/residue pair that best leads to the conformational change of interest, we relate this finding to possible causes. In this study, PRS is applied by scanning each residue in 500 random directions.

To assess the quality of the predicted displacements of all residues resulting from a force applied on selected residue *i*, we use the correlation coefficient between the predicted and experimental displacements, averaged over all the affected residues, *k*:

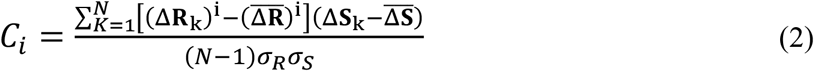

In equation 2, the overbar indicates the average, Δ**S**_k_ are the displacements between the initial and the target forms obtained from the PDB structures, *σ_S_* and *σ_R_* are the respective standard deviations for experimental and calculated displacements. A value close to 1 implies good agreement with the experimental conformational change, while a value of zero indicates lack of correlation between experimental and theoretical findings. Several approaches were taken to select the residues that are effective in directing the protein towards alternative conformations depending on the distribution of the maximum of the *C_i_* values, 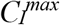, obtained from the 500 perturbations and calculated through eq. 2. We first list 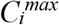 in ascending order: (1) If there is a sharp decrease in the Pearson correlation values, we list the top residues until that gap. (2) If there is a smooth decrease in the Pearson correlation values, we list the residues that are common among top 10 residues of all cut-off values. We also check the location of the residues that do not survive these selection criteria. We have found that the variable residues observed among top residues in different cut-off values are spatial neighbors of the listed ones.

### 2.1. Protein complex selection

We analyzed a set of 25 protein complexes in their bound and unbound forms (Table 1-column 2). We collected protein pair structures from those reported in Benchmark 0.0 of ZDOCK (Chen and Weng, 2002). The complexes are non-redundant and they have X-ray structure solved at better than 2.90 Å resolution. They include a wide variety of function and affinities; they belong to different biological families. The constituent unbound forms of all the 25 complexes are available in the PDB with solution NMR or X-ray structures solved at better than 3.50 Å resolution. For the proteins resolved by solution NMR, we always use the first model for the PRS calculations. More specifically, we have chosen the protein complexes with no less than two missing residues along the protein chain, either in the bound or unbound form. Furthermore, if the number of residues in the bound and unbound components differ, we only analyze the common parts of the bound and unbound proteins.

**Table 1.**
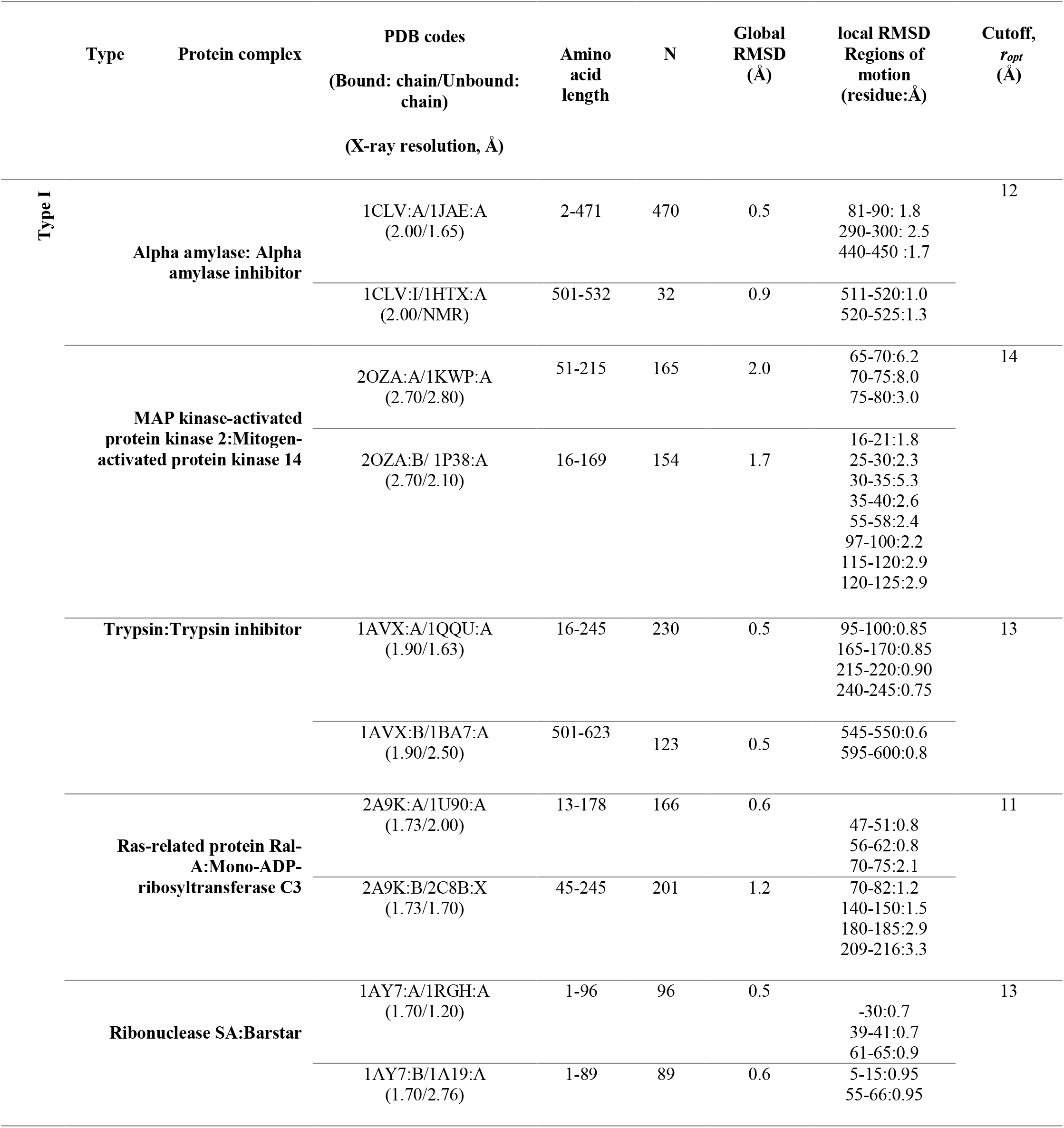
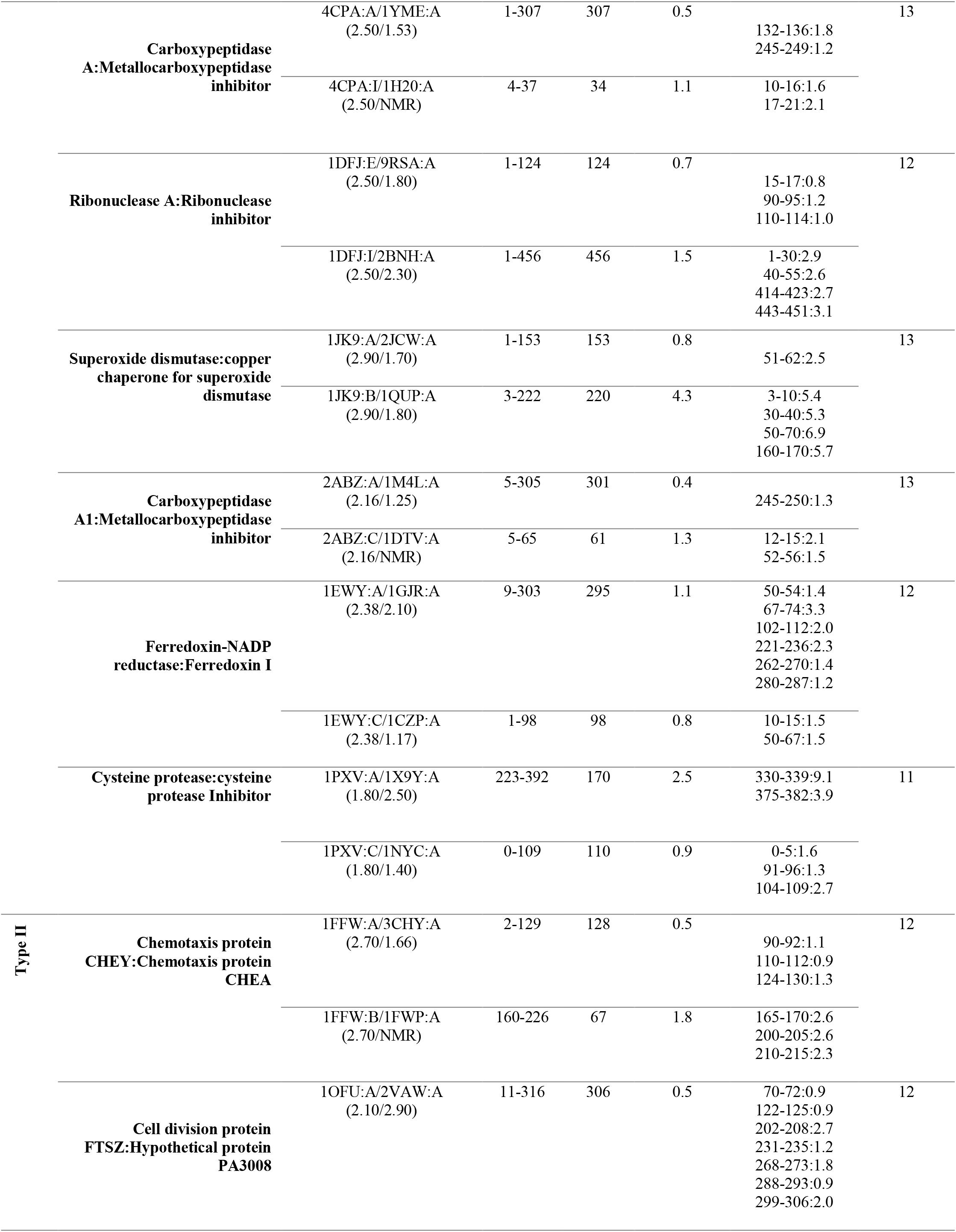
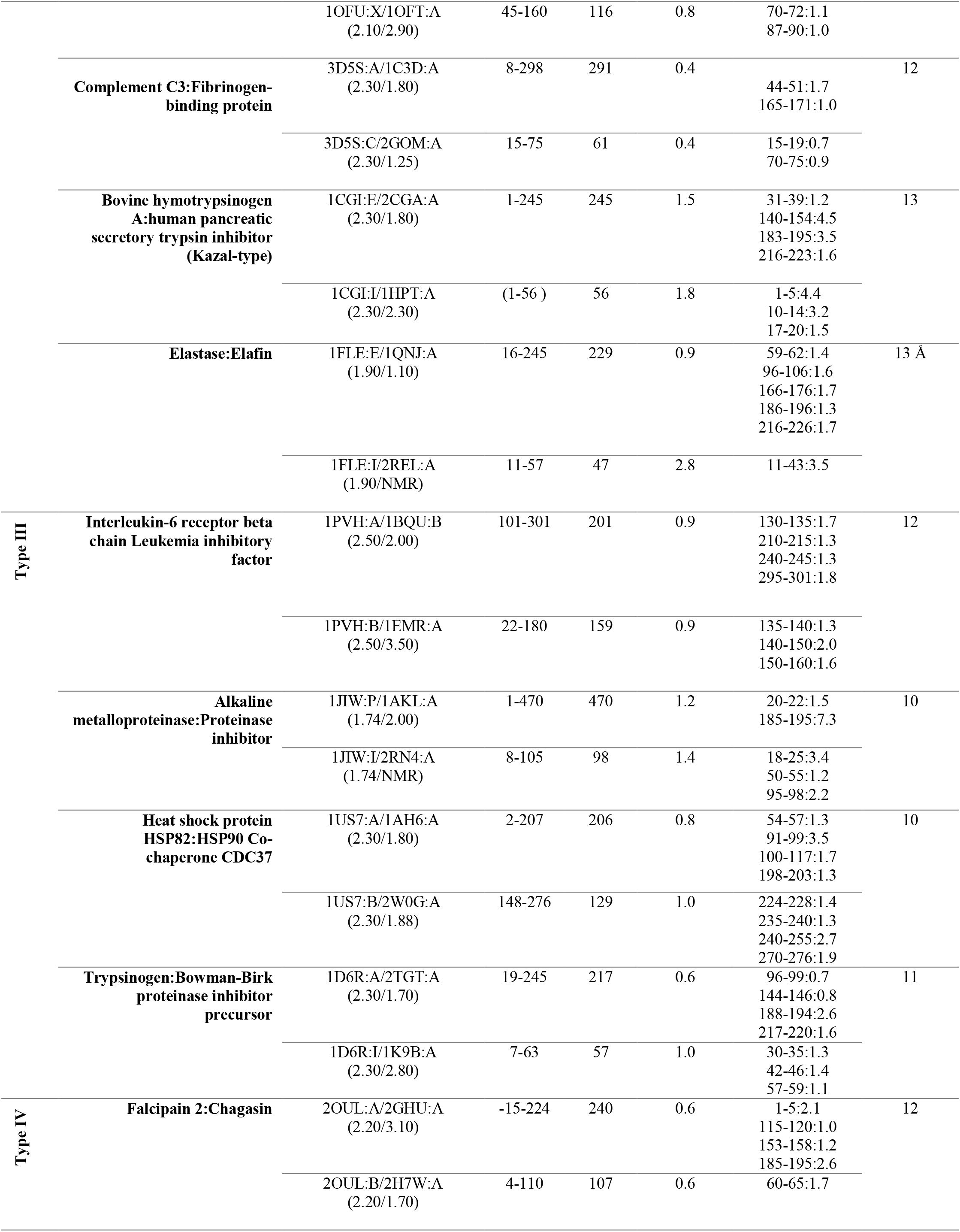
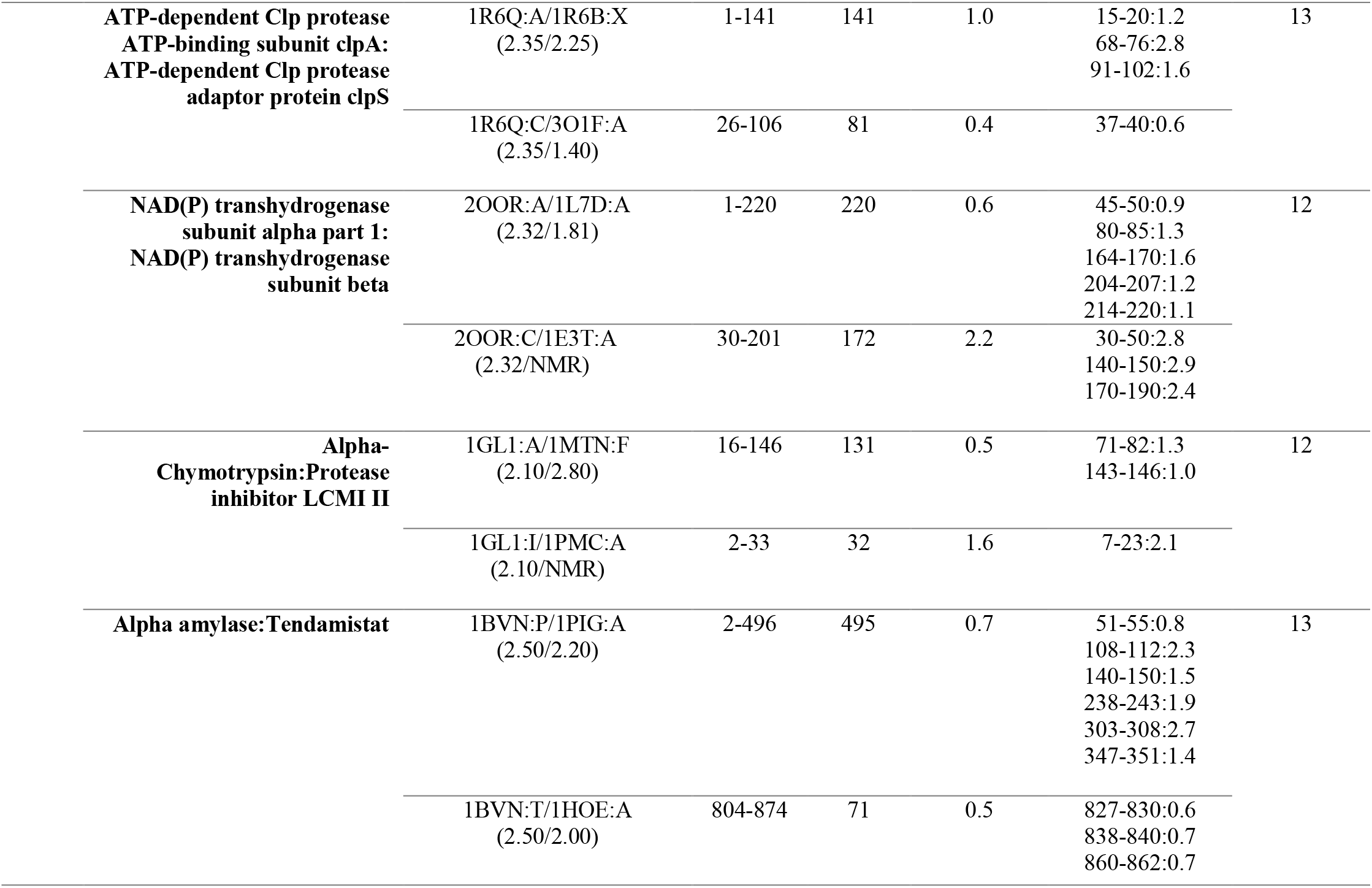
General features of protein complexes studied

### 2.2. Optimization of the cut-off distance

For each pair of experimental structures, the unbound form is superimposed on the bound form, followed by the residue displacement vectors computation, Δ**S**. In this study, we perturb the bound form of each protein by applying a random force to the *C_α_* atom of each residue in the complex. We select residues whose perturbation leads to the Δ**R** vector (equation 1) that best resembles the disassociated proteins using equation 2 as the measure. For a given protein, we select the cut-off distance, *r_opt_* that yields the closest agreement with the displacement vectors from experiments for at least one residue (Table 1. column 8). We verify that the residue indices that provide the best Pearson correlation value are always present within the range perceived as the highest correlation value for all cut-off distances studied. We note that the correlation values reported in Table 2 (column 5 and 6) are not affected for the range of values *r_opt_*±1 Å for all of the 25 protein complexes. Strikingly, although many of these proteins display large Pearson correlation values, the numbers of residues yielding the highest values differ among proteins. For some proteins, there is not much specificity on the residue to be perturbed to reproduce the conformational change. For others, by perturbing a very specific location, the complete conformational change is obtained. The former category is exemplified by fibrinogen-binding protein (pdb code: 3D5S (Haspel et al., 2008)) while serine protease and its proteinaceous inhibitor (pdb code: 1D6R (Koepke et al., 2000)) is an example of a protein complex that we need to perturb in a specific location to mimic the disassociated conformation.

**Table 2.**
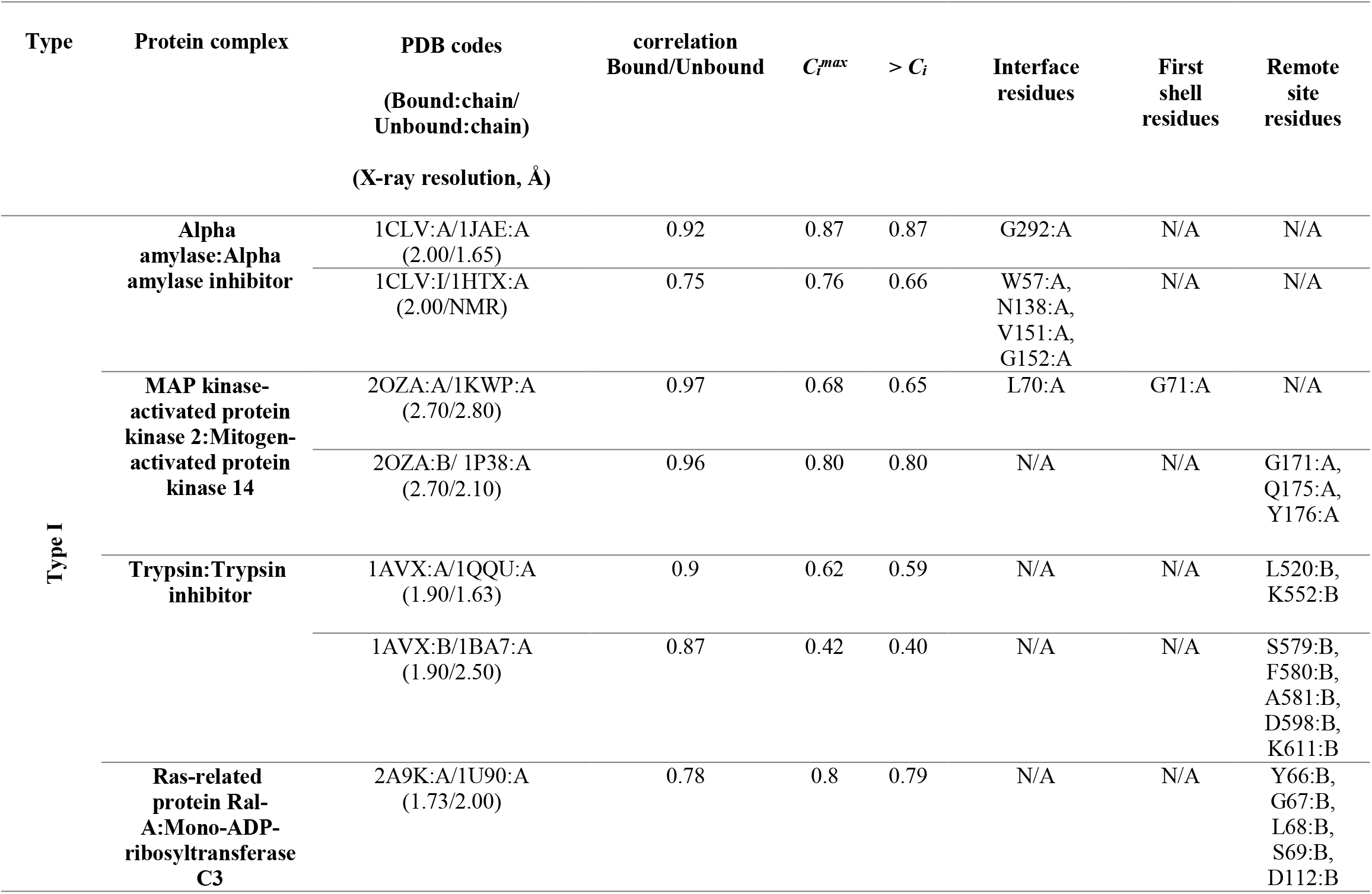
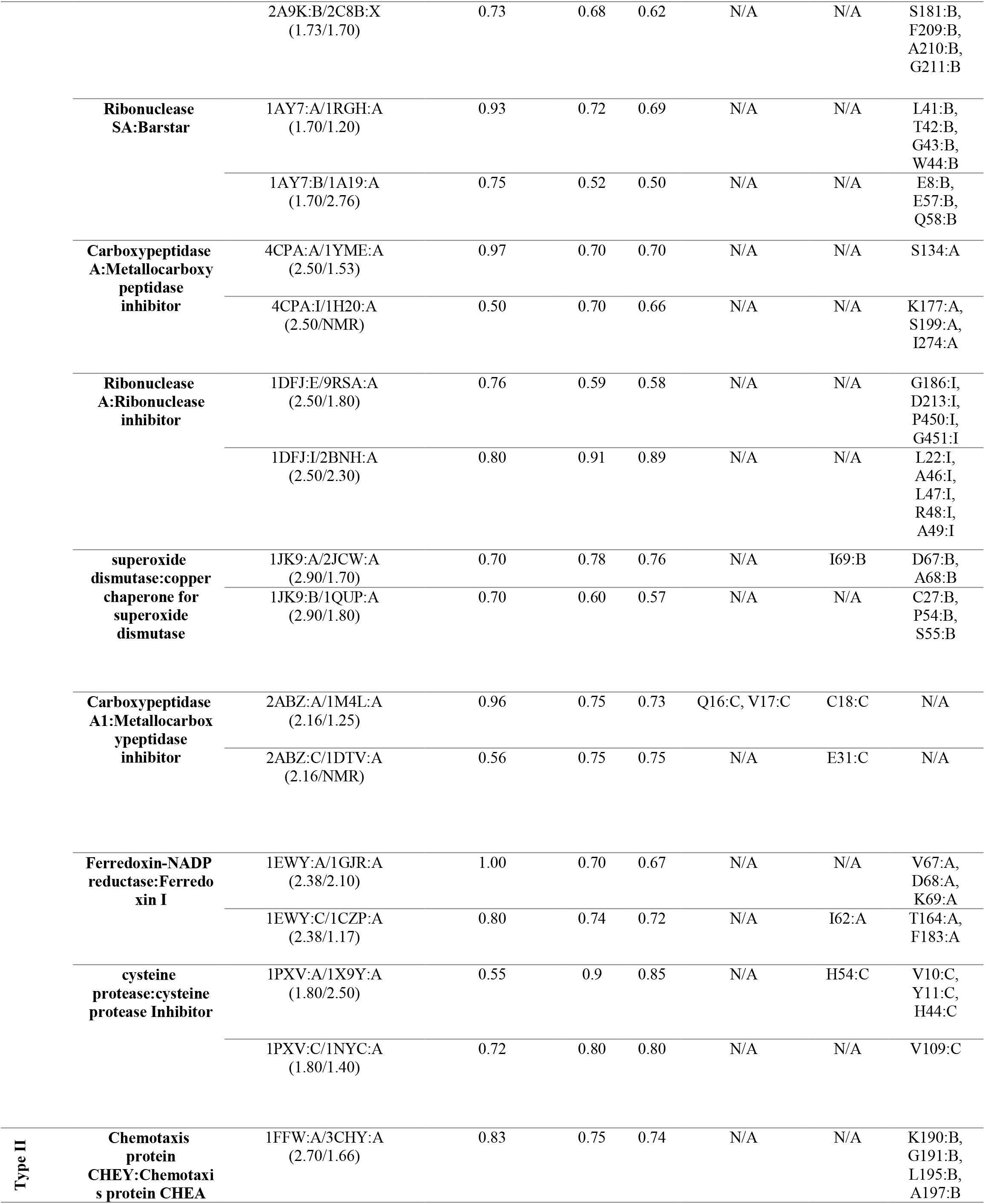
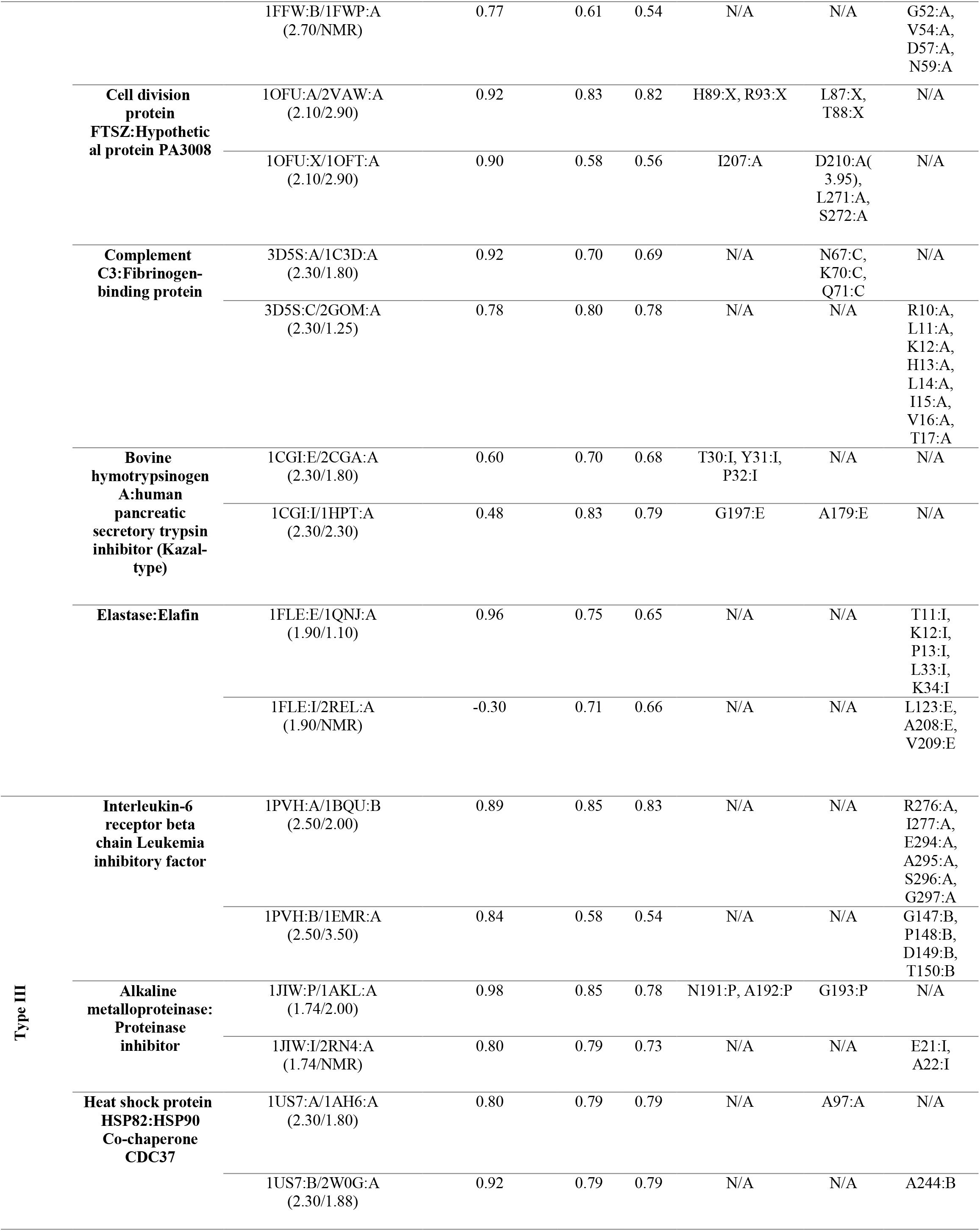
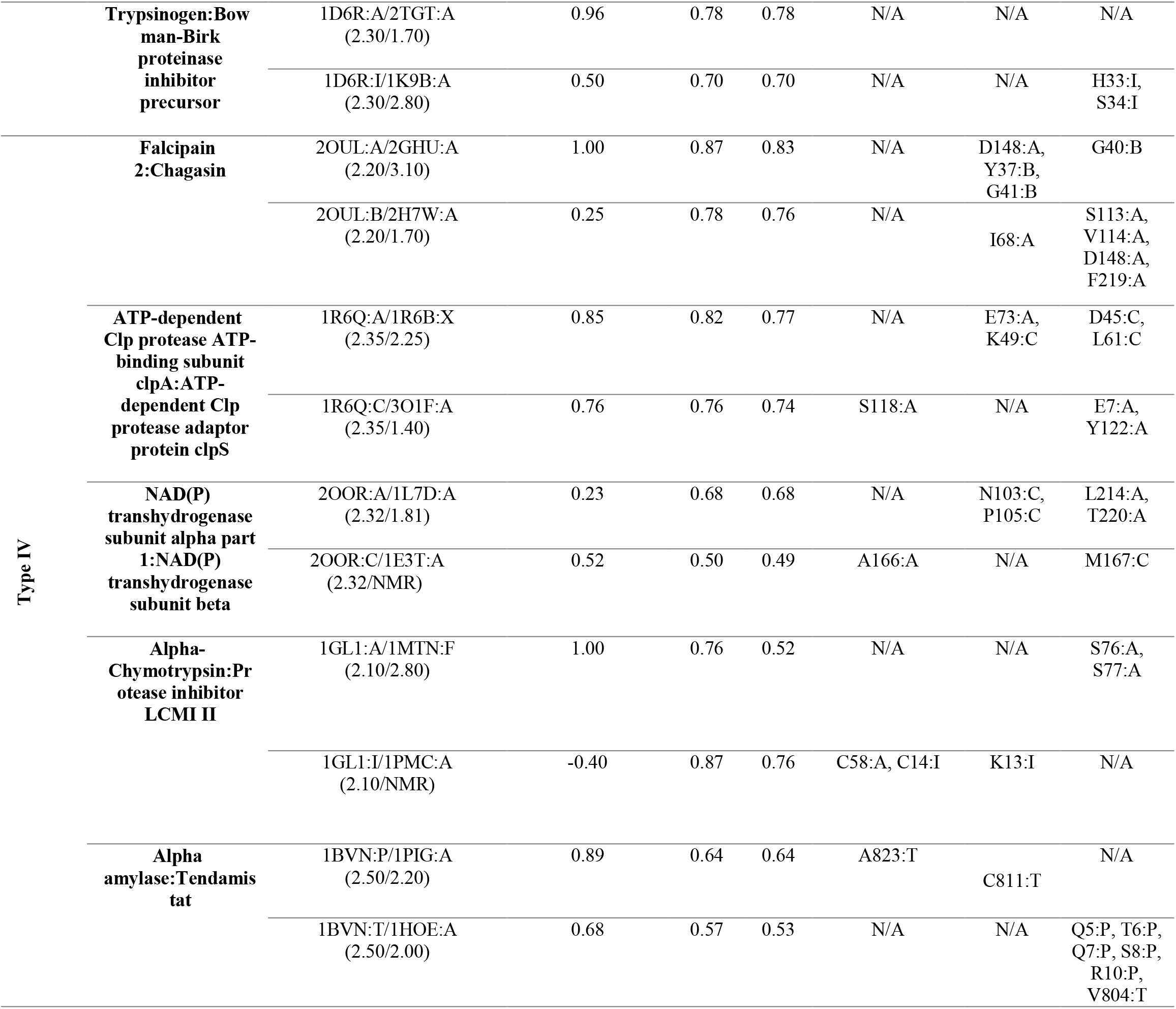
PRS results and classification of the protein complexes

## 3. Results and Discussion

The structural difference between the bound and the unbound forms, based on the Cα superposition of the binding partners, show that while for 22 of the cases the interface RMSD is less than 1.5 Å; for two cases interface RMSD is 1.5-2.2 Å and two cases have interface RMSD greater than 2.2 Å (Humphrey et al., 1996). For the pair of initial and target forms of the proteins present in bound and unbound form, we perform STAMP structural alignment, implemented in VMD 1.9.1 MultiSeq plugin (Humphrey et al., 1996). We record the RMSD between the structures of the target forms with the initial structure which vary between 0.4-4.3Å (Table 1, column 6). Although large motions of side chains and surface loops is always present as a local conformational change, we do not detect any discernible shape change on a global scale in proteins in their bound form compared to the respective unbound constituents. For all cases, the main chains essentially have the same conformation in bound and unbound forms and the binding partners interact as rigid bodies. However, the superposition of mobile segments of the proteins in the complex and corresponding unbound forms produces higher RMSDs (Table 1, column 7) and can be as large as 9.1 Å (see the case of cysteine protease and its inhibitor (Filipek et al., 2003)), the largest deviations being mostly confined to flexible unstructured stretches, i.e. turns and bends.

Recording the residues identified by PRS (Table 2; those with *C_i_* values exceeding the value listed in column 6), displays those encouraging the conformational change from the bound form to unbound form do not necessarily reside on these flexible structures. We have collected 161 effector residues from PRS calculations (Table 2, columns 7-9). While 63 of them are located on flexible loops with large motions and high RMSD values, we identify 30 residues residing on α-helices and 35 on β-strands. The remaining are on loops that do not display any large structural change upon binding.

The average motions of the proteins, quantified by the root mean square fluctuations (RMSFs), are usually expected to dampen upon binding, especially at the binding interface residues, even when the protein conformation is unaltered (Baysal and Atilgan, 2001a). RMSFs of each protein complex constituents in their bound and unbound form are derived from autocorrelation of the residues in each protein pair. By treating the **H^-1^** matrix as an *N*×*N* supermatrix, whose *ij*^th^ element is the 3×3 second moment matrix of correlations between the x-, y-, and z-components of the fluctuations Δ**R**_*i*_ and Δ**R**_*j*_ of residues *i* and j (Baysal and Atilgan, 2001b) are calculated, whose diagonal elements predict the RMSFs (Atilgan et al., 2001). The cut-off distances, *r_opt_*, optimized for building the Hessian matrix of each protein complex have the same values as *r_c_* chosen for PRS analysis. We report the correlation values between proteins in their bound and unbound form for all the protein pairs; the similarity between the RMSF profiles of a protein in its bound and unbound form is expressed as a Pearson correlation and is listed in column 4 of Table 2. We observe that there is a significant change in RMSF of binding region residues in one of the constituents in each pair, so far as to have a negative correlation in some cases; e.g. fluctuation patterns in some regions of the protein is reversed upon complexation. This means that the local fluctuations of the interface area vary in at least one protein upon complex formation, while local fluctuations of their binding partners display the same pattern as the unbound form. However, in an exceptional case of transhydrogenase complex (pdb code: 2OOR (Bhakta et al., 2007)), we observe low correlations, 0.23 and 0.52, between the binding proteins and their unbound form. For this protein complex, the RMSF curves of both the constituents display a significant transformation of fluctuation pattern upon protein binding.

### 3.1. Features of the amino acids involved in disassociation event

From the 8828 residues, 161 of them are selected by PRS. These residues, whose perturbation encourages the unbound over the bound form, are distributed among all amino acid types. The percentage of each amino acid type in our analysis pool and their contributions in PRS analysis are listed in Table 3.

**Table 3.** Types of amino acids selected by PRS compared to all residues in the study

| Residue type | Non-polar |  |  |  |  |  |  |  | Charged |  |  |  |  | Polar Amino |  |  |  |  |  |  |
| --- | --- | --- | --- | --- | --- | --- | --- | --- | --- | --- | --- | --- | --- | --- | --- | --- | --- | --- | --- | --- |
|  | A | V | L | I | M | F | G | W | P | D | E | R | K | N | Q | T | Y | S | H | C |
| PRS residues (%) | 10 | 7 | 9 | 4 | 1 | 3 | 13 | 1 | 3 | 6 | 4 | 3 | 6 | 4 | 4 | 5 | 4 | 9 | 2 | 3 |
| All residues (%) | 8 | 7 | 9 | 5 | 2 | 4 | 8 | 2 | 4 | 6 | 6 | 4 | 6 | 5 | 4 | 6 | 4 | 7 | 2 | 3 |

PRS does not display any preference over amino acid types and any contribution to PRS selection is corroborated to the population of the amino acid type in the total analysis pool.

For example, methionines and tryptophans, each with 1% frequency are the least detected residues by PRS. They are also less frequently seen in the analysis pool (2% of the population). The only residue type that is observed significantly more that in the average pool is glycine which constitutes 13% of all PRS selected residues, compared to its 8 % abundance in the residue data set.

In Table 4, we report the local secondary structure attributes of residues detected by PRS compared to all residues. The secondary structures are assigned by the “Timeline” plugin of VMD and are calculated based on the STRIDE algorithm (Humphrey et al., 1996).

**Table 4.** Secondary structure attributes of amino acids selected by PRS compared to all residues in the study

| | α-helix | π-helix | 3-10 helix | β-strand | Isolated bridge | turn | coil | Total |
| --- | --- | --- | --- | --- | --- | --- | --- | --- |
| **PRS residues (%)** | 19 | 0.0 | 2 | 20 | 1 | 29 | 29 | 100.0 |
| **All residues (%)** | 26 | 0.0 | 4 | 24 | 2 | 26 | 18 | 100.0 |

Among all the protein complexes, we do not find any π-helix type of structure. Residues in the total analysis pool are mostly populated by turns, α-helices and β-strands with 26, 26 and 24% distribution, respectively. However, we find that most preferred regions by PRS defined residues are on coils and turns, each with 29% of all PRS defined residues, although they populate only 44% of the analysis pool. In particular, the enhancement of coil residues in the PRS selection is statistically significant, as these are represented by 29 % in the PRS sub-pool, up from 18 % of all residues in the original pool of residues.

We divide the protein structure into three zones; interface, first coordination and remote, so as to categorize the location of the PRS selected residues. The interface of the two proteins present in the complex is defined by the Cα atoms of the residues from the two sides of the pair residing within 7 Å cut-off distance of each other. We define first coordination shell residues as those located within 7 Å cut-off distance from any interface residue. All remaining residues are classified as remote, defined as those residing beyond the first coordination shell of the interface. We observe that except for the case of alpha amylase and its inhibitor (pdb code:1CLV (Pereira et al., 1999)), PRS selects for remote residues (Table 2). In fact, in 9 cases PRS selected only residues away from the interface. The remaining 16 protein complexes display residues from different parts of the protein in their PRS analysis, including, but not limited to the interface. In fact, these residues are overwhelmingly located on or near the outer surface, as indicated by their depth values from the surface as calculated by the DEPTH server (Tan et al., 2013) and listed in Table S1. In fact, those few that are deeply located (depth greater than 5 Å; shown in bold) are part of a network of interactions whose one end is located on the surface. Thus, the interface of a protein complex is not the controlling region for disassociation of the two proteins.

Remarkably, residues signaling the disassociation of each protein in a given complex are not located on the same protein in all cases. Accordingly, effective sites involved in the disassociation of various protein complexes found in PRS analysis are categorized into four groups based on their responses to the perturbations on the protein. Proteins in which disassociation is signaled through remote residues of the complex are labeled as Type I (11 cases). In this group, PRS top rated residues are all confined to one of the binding proteins. Thus, residues on this protein also control the conformational changes of the binding partner. Type II are the proteins in which residues confined to one of the proteins control the other binding protein and vice versa. Type III are the proteins in which each constituent controls its own disassociation events, and type IV are the proteins where presence of both proteins is essential for the disassociation to occur, as residues signaling the disassociation are scattered on both binding partners. Tables 1 and 2 are organized according to these four distinct groups (I-IV).

### 3.2. Long range control of disassociation is coupled to electrostatic effects

In a subset of the cases, perturbation of specific sites on only one of the constituents in the protein complex modulates disassociation. We label these as Type I group of protein complexes. The functional amino acids defined by PRS which are involved in disintegrating the contact network displays no specific perturbation location in Type I; they may be located on the interface, first shell or remote locations of the protein tertiary structure. Thus, the local perturbations which lead to global conformational shifts between bound and unbound states are not bundled in a specific region.

To determine if these long range effects are controlled by electrostatics, we obtain the electrostatic potential distributions on the biomolecular surface using the APBS package (Baker et al., 2001). In APBS calculations, parameters are set to their default values; i.e., biomolecular and solvent dielectric constant are set to 2 and 78.54, respectively, the radius of the solvent molecules is 1.4 Å and the temperature is 298.15 K; finally, the cubic B-spline discretization method is used to grid biomolecular point charges. Electrostatic effects play a major role in the functionality of this group of protein complexes. Monitoring the electrostatic potential distribution along the protein surfaces in their bound and unbound forms reveals that any given protein in the complex whose electrostatic potential distribution state is stable in their bound and unbound form is also the protein controlling disassociation. Conversely, the binding partner that does not have any PRS determined residue displays considerable change in its charge distribution. In figure 1a, we exemplify how the electrostatic potential distribution on the surface of the protein changes from the free to the bound form.

**Figure 1.**
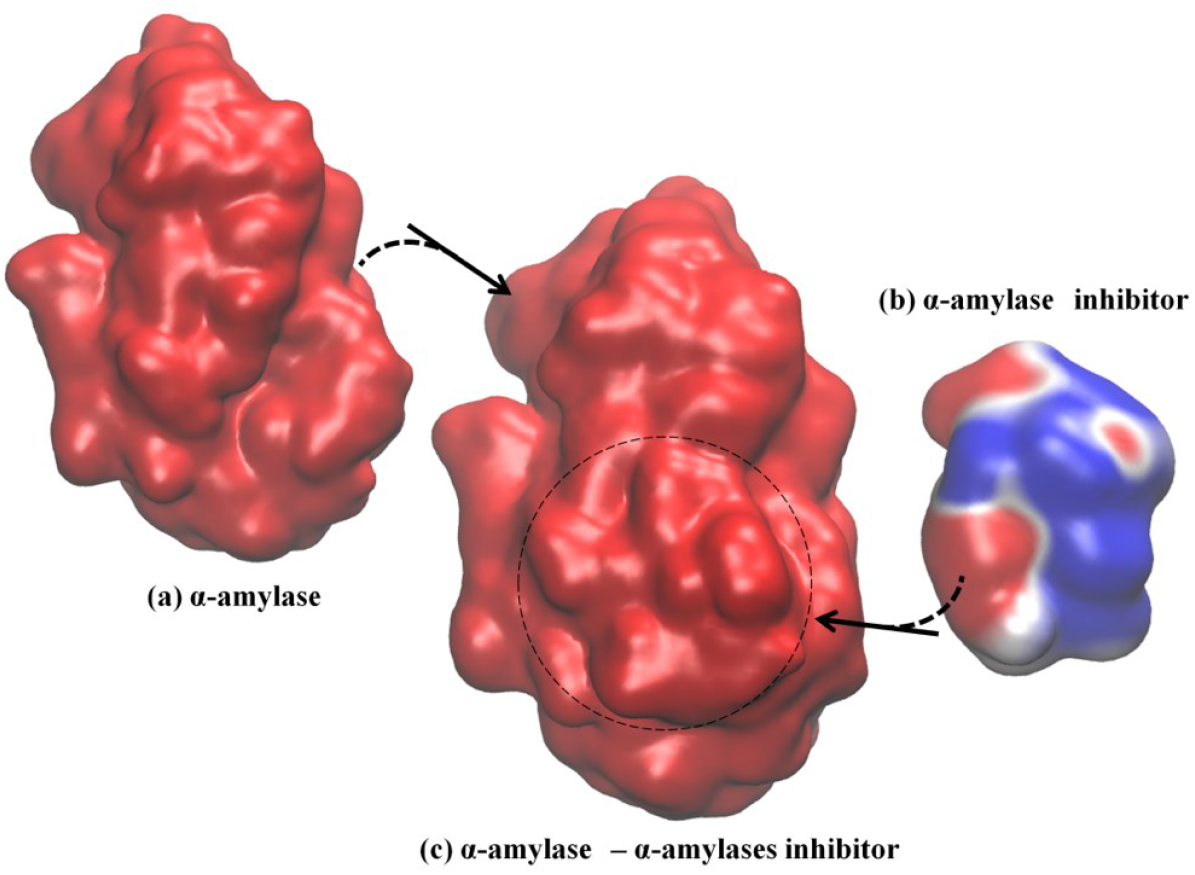
Example of Type I electrostatic isocontour shifts upon binding; α-amylase/α-amylase inhibitor drawn at ±0.5 *k_B_T*/*e*; blue is positive and red is negative. The signaling protein, α-amylase, where all PRS determined residues reside (Table 2), maintains its electrostatic potential distribution while α-amylase inhibitor displays altered electrostatic potential distribution. **(a)** α-amylase in the unbound form, with overall negative electrostatic potential distribution along the surface. **(b)** α-amylase inhibitor in its unbound form with a mixed pattern of negative and positive negative electrostatic potential distribution along the surface. **(c)** α-amylase/α-amylase inhibitor complex with overall negative electrostatic potential distribution. The spatial orientation of the proteins in the complex is kept the same as the respective unbound forms. Dashed circle indicates positioning of the inhibitor in the complex.

Type II group represents another set of protein complexes with remotely controlled conformation changes from the bound to the unbound form. In this group, there is cross-controlled disassociation; i.e., residues that lead to the conformational change upon dissociation on one protein are located on the partner in the complex. We observe that perturbations in a stretch of consecutive resdues is required to trigger the interconversion between two conformational endpoints (see Table 2, Type II). In addition, analysis of the electrostatic potential distribution shows that the proteins interacting with each other possess a similar state of charge distribution. Thus, if the unbound forms had a different electrostatic potential distribution, they reorient themselves to the same state upon protein complex formation. In figure 2, we illustrate electrostatic potential surfaces before and after complex formation in Efb-C and its complement target C3d (pdb code: 3D5S (Haspel et al., 2008)). In fact, it has been reported that formation and stability of Efb-C binding to C3d is electrostatic in nature (Haspel et al., 2008). Kinetic experiments in salt concentrations of 75-600 mM indicate the sensitivity of association/disassociation phases of wild type and various mutants to ionicity (Haspel et al., 2008). This suggests overall electrostatic contribution to be of importance in the initial complex formation and further in stabilizing the complex under the prevailing conditions. Our PRS analysis of Types I and II is further improved by these observations such that under various environmental perturbations, disruption of long-range and short-range electrostatic complementarity seem to impair stability and affect complex formation of binding partners.

**Figure 2.**
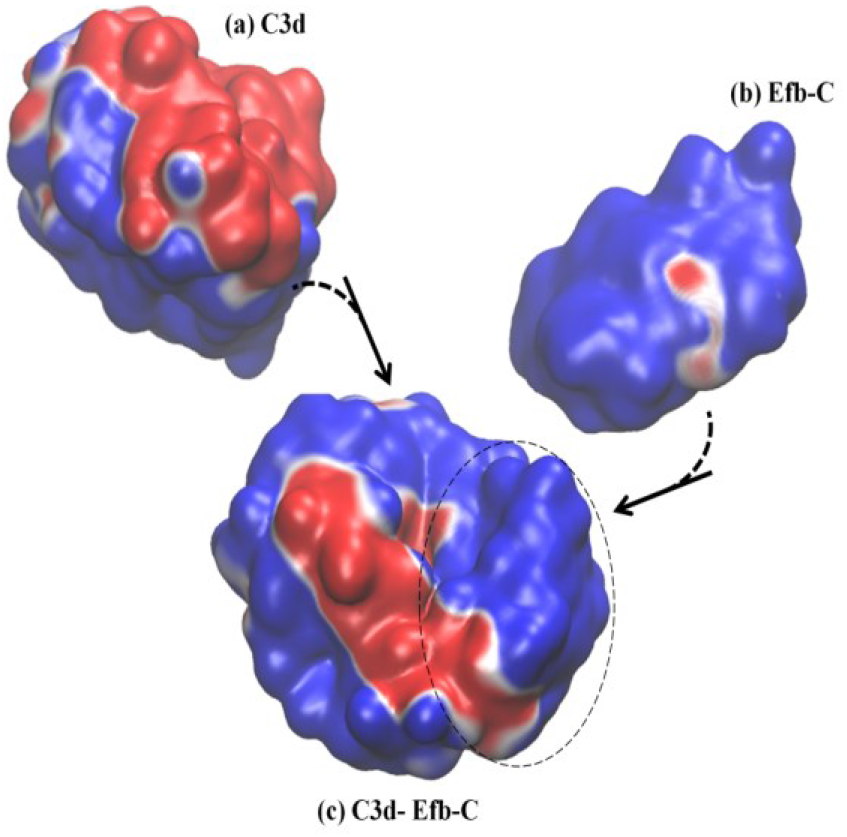
Example of Type II electrostatic isocontour shifts upon binding; Efb-C and its complement target C3d drawn at ±0.5 *k_B_T*/*e*. Blue is positive and red is negative. **(a)** C3d in its unbound form, with a mix of negative/positive electrostatic potential distribution along the surface. **(b)** Efb-C in its unbound form with predominantly positive electrostatic potential distribution along the surface. **(c)** Efb-C/C3d complex. Dashed circle displays Efb-C protein in the complex and the rest of the surface belongs to C3d protein. Both proteins display a mixture of negative/positive electrostatic potential distribution along the surface. In particular, positive surface of the Efb-C displays increased negative areas upon complex formation, while C3d loses negative patches. The spatial orientation of the proteins in the complex is kept the same as that presented in unbound forms.

Remarkably, electrostatic effects provide an excellent description for the observed pattern in both Type I and Type II complexes. Electrostatic interactions are the primary factors of pH dependent processes in biochemical reactions. Particularly, we find that among the 16 protein complexes included into Type I and II groups, 13 of them belong to enzymes. Enzymatic activities are known to be pH dependent and protonation state of catalytic and active site residues are effective in potential distribution of the binding region. Consequently, charge distribution of these regions will modulate the interactions between the proteins and the reaction products. It has been reported that, enzymes make use of their preoriented environment to stabilize the transition state and the reduction in catalytic energy is accomplished by electrostatic stabilization of the active site of the enzyme (Warshel, 1998; Warshel et al., 2006).

In the same vein, we propose that the electrostatic characteristics of residues top-rated by PRS might be found essential to specificity and ligand binding properties in enzymatic reactions. For example, in the Type I complex C3bot-RalA (Ras-related protein Ral-A:Mono-ADP-ribosyltransferase C3, pdb code:2A9K (Pautsch et al., 2005)), displayed in figure 3a, residues 207-214 of C3bot are part of the substrate recognition site and important in catalytic activity. PRS selects residues 209-211as candidate positions to provoke the disassociation of the complex. Another key region involved in disassociation upon force application is residue 112 of C3bot, which also contributes to complex formation. In addition, residue 181 is located on a loop responsible in substrate recognition. Accordingly, perturbation of specific sites found by PRS analysis and experimentally verified to be functionally important might reorient the enzyme/substrate dipoles that organize the catalysis and destabilize the charged transition state (Thomas et al., 1985). These changes might thus prevent the binding event or promote disassociation after the chemical reaction has terminated.

**Figure 3.**
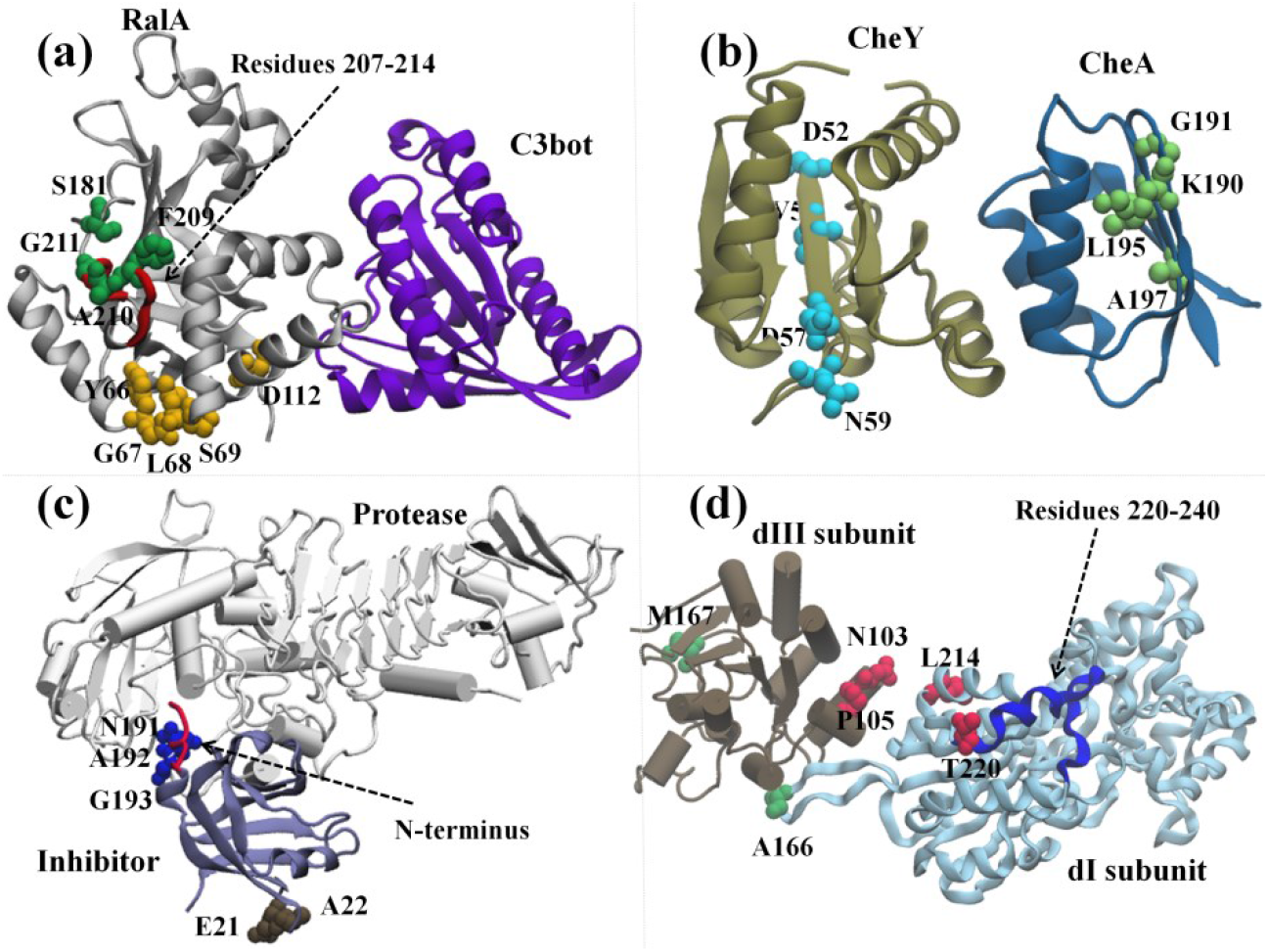
Example complexes demonstrating dissociation scenarios discussed: **(a)** Ribbon representation of C3bot-RalA complex as an example of Type I proteins; C3bot is in purple and RalA is in silver. PRS selected residues that control the disassociation of the complex are displayed as spheres and all are confined to RalA compartment. Residues controlling the disassociation of RalA are in yellow, while the residues controlling disassociation of C3bot are in green. Substrate recognition site of C3bot (Residues 207-214) is shown in Red. **(b)** Ribbon representation of chemotaxis CheA:CheY complex as an example of Type II proteins; CheY is in tan and CheA is in blue. Residues controlling the disassociation of CheY are located on CheA and are displayed in green while residues controlling the disassociation of CheA are located on CheY and are displayed in cyan. **(c)** Ribbon representation of alkaline protease and its cognate inhibitor as an example of Type III proteins. Protease is represented as white cartoon and the inhibitor is in ice blue. Residues controlling the disassociation of protease are located on it and displayed in blue. **(d)** Ribbon representation of dI2dIII1complex of proton-translocating transhydrogenase as an example of Type IV proteins. Residues that control the disassociation of the dI subdomain are located on both subunits and are displayed in red. Residues that control the disassociation of dIII subunit are displayed in green.

We also find that the effect of the local charge distribution on enzyme functions is not limited to the active site and that the remote locations on the protein are effectively involved in the disassociation process. Mutations of charged patches remote from either the protein or ligand binding site might alter the binding kinetic rates, shift pK_as_ and lead to weak molecular recognition (Thomas et al., 1985). In response to a particular perturbation, exposure to a different environment, reorganization of charged atomic groups and dielectric relaxation of the protein affects the electrostatic potential distribution of the interface or active site region considerably, creating the steering forces that guide the disassociation reaction. Thus, a local change of interactions at a remote site leads to a global structural change that modifies the organization of the interface contact network and leads to disassociation of the two proteins. The contribution of this distal perturbation on enzyme/inhibitor activity basically may be viewed as leading to a cooperative conformational transition. In both Type I and Type II, the information transmission between remote functional sites on one protein and the entire structure of the interacting partner naturally occurs via the interface linkage. Iin such proteins, the network of contacts in the interface could form a so-called “conductive” region so that the signal from one protein is transferred through the interface to control the functionality of the second binding protein.

As one of the test beds, we have studied chemotaxis CheA:CheY complex (pdb code: 1FFW (Gouet et al., 2001)). PRS finds residues 52, 54, 57 and 59 of CheY to play a role in selecting the unbound over bound conformation in the presence of an external force (figure 3b). This complex is an example of remote communication between a two-component signal-transducer. pH dependent catalytic activity has also been observed in chemotaxis CheA:CheY, in which ligand binding on CheA is conducted to CheY. Phosphorylation reaction on H48 of CheA subsequently transfers to D57 of CheY and the complex disassociates (Silversmith et al., 1997). CheY itself is incapable of providing an acidic residue during the phosphorylation event and the complex formation with CheA results in a conformational change on CheY as an acidic residue (D57) donates a proton to a phosphodonor in an optimal orientation and the protein-protein phosphotransfer occurs (Silversmith et al., 1997). The pH dependence of the phosphotransfer kinetics in the pH range of 7.5-10, studied through two mutants of CheY active site residues shows simply a moderate decrease in rate constants compared to the wild type CheY. This observation suggests that conserved active site residues do not have an essential and direct role in catalysis. Thus, the loss of activity throughout this range for phosphotransfer to wild type CheY is attributed not to deprotonation events on CheY; rather it is likely due to deprotonations in CheA (Silversmith et al., 1997). Variability of position 59 of CheY as a non-conserved and indirect active site residue in modulating the autophosphorylation of CheY with small molecule phosphodonors shows no detectable binding between the phosphodonor and CheY, validating the significant impact of position 59 on autophosphorylation kinetics (Thomas et al., 2013). Mutation of N59 to R, K, M, L, A, D and E results in both increase and decrease in autophosphorylation rate constants. Substitution with positively charged residues increases the kinetic rates while substitution with negatively charged residues decreases the rate, implying how local electrostatic interactions at position 59 modulate the CheY autophosphorylation (Thomas et al., 2013).

### 3.3. Auto-controlled conformational transitions are correlated to mechanical organization of the protein in Type III complexes

In Type III group of protein complexes, disassociations of either constituent of the complexes are governed by a local perturbation on the respective protein. We label this behavior as auto-controlled disassociation. In this category, all the residues involved in global transformation of bound to unbound forms are located out of the interface region, trypsinogen and its Bowman-Birk proteinase inhibitor precursor (pdb code: 1D6R (Koepke et al., 2000)) being an exception. We observe that in Type III complexes, the global RMSD between bound and unbound forms of each protein varies between 0.6-1.4 Å. However, regions of high mobility exist in which the local RMSD may take values as high as 7.3 Å (Table 1). Such regions belong to unstructured surface exposed loops where they display large deviations upon complex formation. The amino acids identified by PRS in this group belong to these regions and very specific locations may be perturbed so that the local fluctuations of the amino acids choose the conformational switch to the target structure. Thus, mechanical motions of the loops produce essential conformational transitions under such point perturbations.

Contrary to Type I and II complexes, the electrostatic potential distribution along the protein surfaces of Type III reveals that each protein maintains the same electrostatic potential distribution state in their respective bound and unbound forms. This means that for any protein complex in this group, orientation of the negatively and positively charged surfaces of the constituents has the same distribution in their respective free forms. Thus, the analysis identifies an “insulating” interface area that prevents the allosteric communication between interacting proteins and each protein functions independently under various perturbations. We note that while there are also stretches with large RMSDs in Type II protein complexes, those regions cross-control the disassociation of the binding proteins remotely and their own shape changes are not mechanically controlled.

As an example case, for the alkaline protease and its cognate inhibitor (pdb code: 1JIW (Hege et al., 2001)) classified as Type III, PRS finds residues 191-193 of protease which are in direct contact with the N-terminus of the inhibitor (figure 3c). The latter has been shown to coordinate the catalytic zinc anion (Hege et al., 2001). Such an interaction adds to the structural stability and leads to a low disassociation constant. Upon any kind of perturbation applied to residues 191-193 of the protease, disruption of the interactions in this region would change the extended conformation of the N-terminus and modify the zinc coordination and thus might facilitate the disassociation process.

### 3.4. Combined perturbations on both partners is essential for disassociation in Type IV complexes

In Type IV group of protein complexes, disassociation of the two constituents might be triggered by point perturbations on either subunit. This contrasts Type III complexes where perturbation of any subunit will mediate the disruption of interactions in the protein pair, facilitating disassociation. Different functional subdomains may exist on these proteins which contribute to binding to diverse set of ligands in their functional pathway and promote disassociation. Additionally, complexes in our test set involving transmembrane proteins also fall into this group. Presence of each fragment of the protein pair in different compartments of the cell environment assist their exposure to different perturbation scenarios and support the idea of simultaneous perturbation of both partners of the protein complex in the disassociation process. As an example, we focus on dI2dIII1complex of proton-translocating transhydrogenase (pdb code:2OOR (Bhakta et al., 2007)). The complex is found in the membrane compartment of the bacteria or animal cells. Proton transfer across the membranes is facilitated by conformational changes of transhydrogenase. PRS identifies residues A166, L214 and T220 of dI subunit and residues N103, P105 and M167 of dIII subunit to be effective in disassociation of the complex (figure 3d). dI and dIII protrude from the membrane while a third compartment dII spans the membrane. Thus, each part of the protein is exposed to a different part of the cell, making the protein susceptible to alternative perturbation scenarios. Residue T220 of dI is part of a loop (residues 220-240) that becomes less mobile upon ligand binding due to surface closing of the protein (Bhakta et al., 2007). Furthermore, M167 of dIII is in the neighboring site of H2NADH and any perturbation on this site might alter the proton pump reactions due to changes made to interaction network of ligand binding region. Accordingly, any perturbation on H2NADH binding region in dIII subdomain may alter the structural features of dI subdomain through remote communication.

## 4. Conclusions

There is plethora of work addressing association of proteins partaking in complexes, and the consensus is to focus on the interface to determine the major features of binding events (Gainza et al., 2020), concentrating on, e.g., pockets formed upon complexation (Li et al., 2004), prediction of binding energies based on the interface (Moreira et al., 2007), close-range electrostatic interactions (Kumar and Nussinov, 2002), and conserved residues along the interface (Kumar and Nussinov, 2002; Li et al., 2004). However, to alter protein functions, e.g. for therapeutic applications, it is also essential to understand the mechanisms affecting their dissociation, a question that has not been thoroughly explored, to the best of our knowledge. In this work, we have studied the characteristics of residues responsible for the dissociation of a set of 25 non-redundant protein complexes, using PRS as the predictor of residues whose perturbation encourages the unbound conformations. Significance of the residues identified by PRS are discussed in detail for four sample cases (Figure 3).

In a statistical analysis of the PRS identified residues, we find that in terms of residue types, the only significant enhancement is in glycine residues, up from 8 % of all residues found in the protein set to 13 % in the subset of residues implicated in protein dissociation. This is in contrast to the studies reporting on hotspots on the interaction surface, whereby tryptophan (21%), arginine (13.3%), and tyrosine (12.3%) have the highest probabilities of occurrence (Moreira et al., 2007). Moreover, PRS identifies residues labelled as controlling dissociation are also significantly enhanced on loops, and are predominantly located on the complex surface, remote from the interface. This finding is plausible, since in contrast to an association event whereby interface compatibility is a major determinant, exposed residues are expected to partake in disrupting the complex.

We find that dissociation events disclosed by PRS analysis may be classified into four main groups as summarized in Table 5.

**Table 5.**
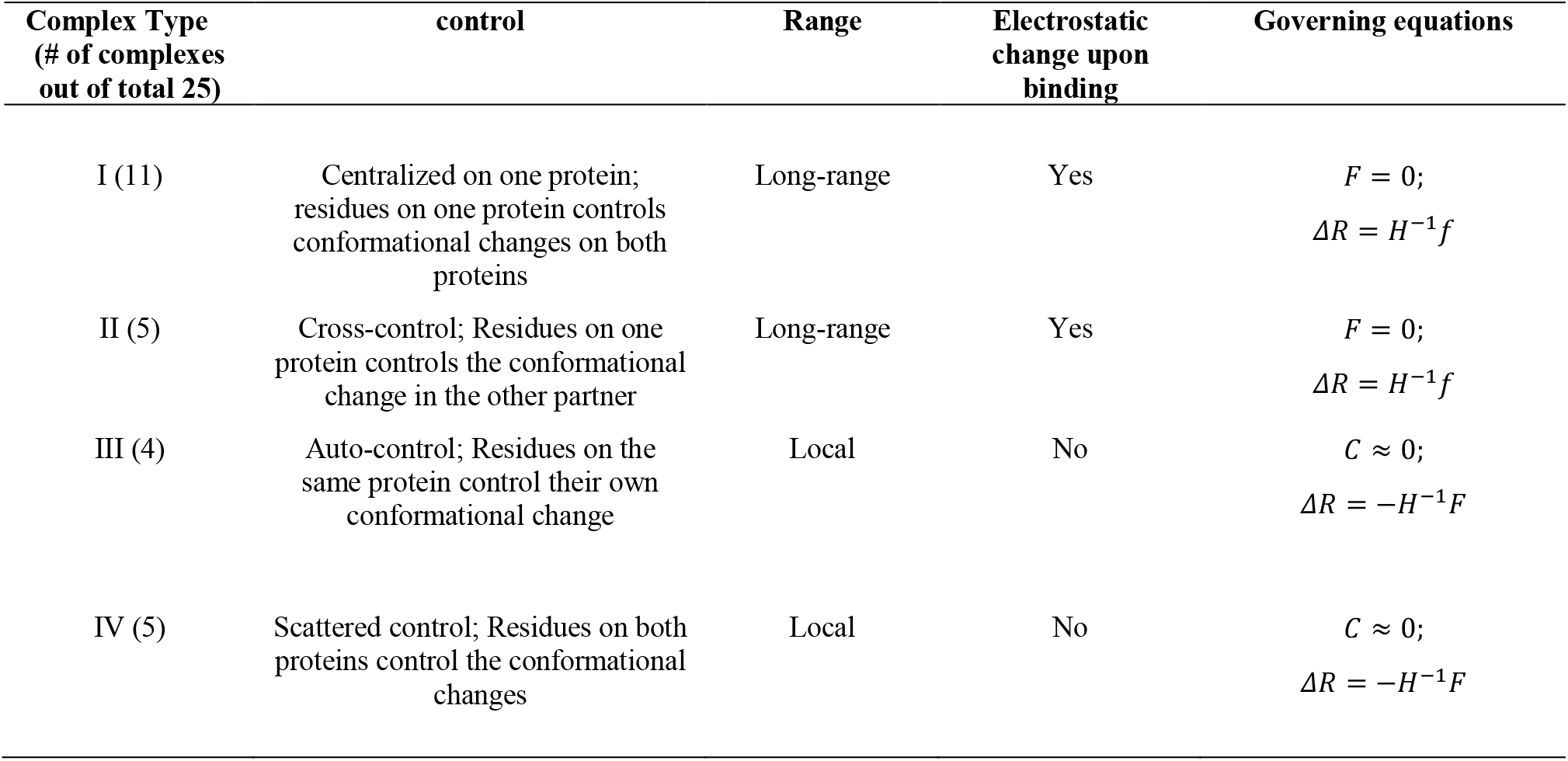
Summary of types of complexes and properties of their dissociation mechanisms

| Complex Type<br>(# of complexes<br>out of total 25) | control | Range | Electrostatic<br>change upon<br>binding | Governing equations |
| --- | --- | --- | --- | --- |
| I (11) | Centralized on one protein;<br>residues on one protein controls<br>conformational changes on both<br>proteins | Long-range | Yes | $F = 0;$<br>$\Delta R = H^{-1}f$ |
| II (5) | Cross-control; Residues on one<br>protein controls the conformational<br>change in the other partner | Long-range | Yes | $F = 0;$<br>$\Delta R = H^{-1}f$ |
| III (4) | Auto-control; Residues on the<br>same protein control their own<br>conformational change | Local | No | $C \approx 0;$<br>$\Delta R = -H^{-1}F$ |
| IV (5) | Scattered control; Residues on both<br>proteins control the conformational<br>changes | Local | No | $C \approx 0;$<br>$\Delta R = -H^{-1}F$ |

The nature of the events leading to dissociation are either expected to be due to mechanical perturbations arriving at certain locations on the surface, or due to environmental triggers that interfere with the electrostatic potential distribution of the complex. In the latter case, a signature event is in the change of the electrostatic potential distributions of one of the binding partners (see Type I in tables 1 and 2; and figure 1) or both of them (Type II complexes; figure 2). The physics of these observations are resolved by an electro-mechanical coupled elastic network model proposed by our group (Sensoy et al., 2017).

Accordingly, in the general case where mechanical motions are coupled to the electrostatics, the potential function *V* in an elastic network may be expressed as,

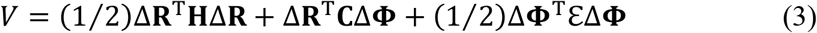

where **ΔR** is the vector of dimension 3*N* for displacements of residues from their average position, **H** is the 3*N* × 3*N* Hessian matrix, Δ**Φ** is carries information on the change in the electric field at the location of each residue due to variations in the environment; the *N × N* diagonal matrix 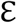 represents permittivity at each position, while **C** matrix represents the coupling between mechanical motions and electrostatics. Two constitutive relations for the force applied, **F**, and the electric displacement **D** are obtained from the derivatives of equation 3 with respect to position and electrostatic potential, 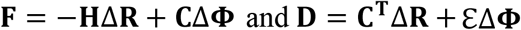. By substitution and rearrangement, we get,

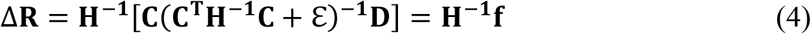

where the term in square brackets is an equivalent force arising from the change in the electric displacement, e.g. due to pH or ionic strength variations in the environment. Thus, we find that even in the absence of an external force, positional displacements may still be obtained, provided there is electromechanical coupling.

The role of PRS is to identify residues where such equivalent forces are focused on and are relevant to the observed conformational change. Conversely, in the absence of coupling, *C* = 0, equation 4 is reduced to the usual expression for a system of springs, i.e. that used in elastic network models; Δ**R** = −**H**^−1^**F**. Under such circumstances, the conformational change may indeed be triggered by an external force, e.g. upon binding of a ligand. Since both equations are of the same form and dimensionality, PRS either identifies the residues that facilitate displacements by mechanical perturbations (Type III and IV), or those which are mechanical mimics to the effects expedited by changes in electric displacement (Type I and II).

This study is a step towards developing descriptors aimed at disrupting protein complexes with the aim of developing therapeutic approaches to alter the function of proteins working in tandem. In particular, targeting remote sites to destabilize interacting proteins using unique approaches will aid in the emerging field of allosteric drug design (Guarnera and Berezovsky, 2020).

## Acknowledgments

This work was supported by the Scientific and Technological Research Council of Turkey (grant number 116F229).

