## Supplementary Information for "A coarse-grained methodology identifies intrinsic mechanisms that dissociate interacting protein pairs"

**Table S1. Depths of residues singled out by PRS in the complex and unbound structures.**

| Type | Complex | PRS residue depth (Bound/Unbound) (Å) | | | | | | | | | | |
|---|---|---|---|---|---|---|---|---|---|---|---|---|
| I | | | | | | | | | | | | |
| | **1CLV(AI)** | **W57:A(10.5/ 6.6)** | N138:A(4.7/ 4.2) | **V151:A(8.1/ 3.8)** | **G152:A(5.5/ 3.6)** | G292:A(3.9/ 4.7) | | | | | | |
| | **2OZA(AB)** | **L70:A(3.9/ 5.9)** | G71:A(3.4/ 3.3) | **G171:A(7.6/ 8.0)** | **Q175:A(5.0/ 5.0)** | Y176:A(3.9/ 4.3) | | | | | | |
| | **1AVX(AB)** | **L520:B(5.7/ 5.5)** | K552:B(4.9/ 4.9) | S579:B(4.0/ 3.7) | F580:B(3.8/ 3.5) | A581:B(3.5/ 4.1) | D598:B(3.6/ 3.4) | K611:B(3.1/ 3.5) | | | | |
| | **2A9K(AB)** | Y66:B(4.6/ 4.4) | G67:B(3.1/ 3.1) | L68:B(4.8/ 4.9) | S69:B(3.5/ 3.6) | **D112:B(8.1/ 4.2)** | S181:B(4.0/ 3.6) | F209:B(4.1/ 3.9) | A210:B(3.9/ 4.8) | G211:B(3.5/ 3.3) | | |
| | **1AY7(AB)** | E8:B(4.8/ 4.2) | **L41:B(6.6/ 6.1)** | T42:B(4.0/ 3.9) | G43:B(3.8/ 3.3) | W44:B(3.7/ 4.0) | **E57:B(4.1/ 5.3)** | **Q58:B(4.6/ 5.6)** | | | | |
| | **4CPA(AI)** | S134:A(3.7/ 3.7) | **K177:A(5.2/ 5.1)** | **S199:A(6.1/ 4.4)** | I274:A(4.8/ 4.4) | | | | | | | |
| | **1DFJ(EI)** | **L22:I(6.8/ 7.8)** | A46:I(4.7/ 4.6) | **L47:I(7.7/ 7.4)** | R48:I(4.9/ 4.7) | A49:I(3.9/ 3.6) | **G186:I(6.4/ 7.1)** | D213:I(4.2/ 4.2) | P450:I(4.2/ 3.9) | G451:I(3.3/ 3.4) | | |
| | **1JK9(AB)** | C27:B(3.8/ 3.9) | **P54:B(6.5/ 6.6)** | S55:B(4.5/ 4.4) | D67:B(4.4/ 4.0) | **A68:B(5.6/ 5.7)** | I69:B(4.3/ 5.4) | | | | | |
| | **2ABZ(AC)** | **Q16:C(6.9/ 5.6)** | **V17:C(8.6/ 7.1)** | **C18:C(6.6/ 5.8)** | E31:C(4.8/ 3.5) | | | | | | | |
| | **1EWY(AC)** | **I62:A(8.9/ 9.7)** | V67:A(4.2/ 4.5) | D68:A(4.3/ 4.3) | K69:A(3.9/ 3.5) | **T164:A(11.3/ 11.9)** | **F183:A(5.8/ 6.9)** | | | | | |
| | **1PXV(AC)** | **V10:C(5.2/ 5.1)** | Y11:C(4.1/ 3.6) | H44:C(4.4/ 4.6) | **H54:C(4.7/ 5.3)** | | | | | | | |
| II | **1FFW(AB)** | G52: A (3. 9/ 3.7) | **V54:A(10.2/10.0)** | **D57:A(9.0/7.6)** | N59:A(4.5/4.4) | K190:B(4.2/4.4) | G191:B(3.3/3.5) | **L195:B(5.7/4.5)** | **A197:B(7.1/ 7.4)** | | | |
| | **1OFU(AX)** | **I207:A(6.8/4.4)** | **D210:A(4.0/6.7)** | **L271:A(7.0/5.3)** | S272:A(4.6/4.0) | **L87:X(3.9/5.5)** | T88:X(4.5/4.0) | **H89:X(6.8/4.2)** | **R93:X(6.2/7.1)** | | | |
| | **3D5S(AC)** | **R10:A(6.5/4.4)** | **L11:A(9.0/6.1)** | **K12:A(6.7/4.4)** | **H13:A(7.3/3.5)** | **L14:A(8.1/5.4)** | **I15:A(6.6/4.9)** | **V16:A(6.7/4.1)** | **T17:A(5.8/4.1)** | N67:C(4.7/4.7) | K70:C(3.6/4.2) | Q71:C(3.6/3.9) |
| | **1CGI(EI)** | A179:E(4.5/4.1) | **G197:E(14.6/9.7)** | T30:I(4.3/4.4) | **Y31:I(5.6/4.7)** | **P32:I(7.2/4.3)** | | | | | | |
| | **1FLE(EI)** | L123:E(4.3/4.4) | **A208:E(5.2/5.6)** | **V209:E(7.2/8.0)** | T11:I(3.3/4.0) | K12:I(4.3/3.5) | P13:I(3.6/3.8) | L33:I(4.4/4.4) | K34:I(3.9/4.3) | | | |
| III | **1PHV(AB)** | **R276:A(7.5/ 7.6)** | **I277:A(9.1/ 9.1)** | E294:A(4.1/ 4.2) | A295:A(3.9/ 3.7) | S296:A(4.3/ 3.7) | G297:A(4.1/ 3.8) | G147:B(4.8/ 4.1) | P148:B(3.5/ 3.7) | D149:B(3.8/ 4.0) | T150:B(3.2/ 4.4) | |
| | **1JIW(PI)** | N191:P(4.6/ 3.6) | **A192:P(5.4/ 3.5)** | G193:P(3.4/ 3.5) | E21:I(3.8/ 4.0) | A22:I(3.6/ 3.7) | | | | | | |
| | **1US7(AB)** | A97:A(4.3/ 4.1) | A244:B(4.1/ 3.4) | | | | | | | | | |
| | **1D6R(AI)** | **G193:A(6.6/ 5.8)** | H33:I(4.4/ 4.1) | S34:I(3.6/ 3.5) | | | | | | | | |
| IV | **2OUL(AB)** | **I68:A(7.3/7.0)** | S113:A(4.4/4.6) | **V114:A(5.5/5.4)** | D148:A(3.9/4.7) | **F219:A(7.8/7.7)** | **Y37:B(7.5/7.6)** | G41:B(4.4/4.4) | | | | |
| | **1R6Q(AC)** | **E7:A(7.0/4.9)** | E73:A(3.9/3.6) | **S118:A(6.9/3.8)** | **Y122:A(6.2/4.2)** | D45:C(4.1/ 4.2) | K49:C(4.5/3.8) | **L61:C(4.7/ 5.2)** | | | | |
| | **2OOR(AC)** | **A166:A(5.6/ 3.6)** | L214:A(4.2/ 3.9) | T220:A(4.3/ 4.2) | N103:C(4.0/ 3.8) | P105:C(3.9/ 4.1) | M167:C(4.0/3.5) | | | | | |
| | **1GL1(AI)** | **C58:A(5.3/5.4)** | S76:A(3.6/ 3.4) | S77:A(3.7/3.9) | K13:I(3.8/3.6) | C14:I(3.9/3.8) | | | | | | |
| | **1BVN(PT)** | Q5:P(4.1/ 4.3) | **T6:P(4.9/ 5.0)** | Q7:P(3.8/ 4.3) | S8:P(3.5/ 3.8) | **R10:P(4.9/ 5.2)** | V804:T(3.4/ 3.7) | C811:T(3.7/ 3.8) | **A823:T(8.5/ 7.3)** | | | |
